# B cells adapt their nuclear morphology to organize the immune synapse and help antigen extraction

**DOI:** 10.1101/2021.04.20.440571

**Authors:** Romina Ulloa, Oreste Corrales, Fernanda Cabrera, Jorge Jara-Wilde, Juan José Saez, Christopher Rivas, Jonathan Lagos, Steffen Härtel, Clara Quiroga, Edgar R Gomes, María-Isabel Yuseff, Jheimmy Diaz Muñoz

## Abstract

Upon interaction with immobilized antigens B cells form an immune synapse, where actin remodeling and re-positioning of the microtubule-organizing center (MTOC) together with lysosomes can facilitate antigen extraction. B cells have restricted cytoplasmic space, mainly occupied by a large nucleus, yet the role of nuclear morphology in the formation of the immune synapse has not been addressed. Here we show that, upon activation, B cells re-orientate and adapt the size of their nuclear groove facing the immune synapse, where the MTOC sits and lysosomes accumulate. Silencing nuclear envelope proteins, Nesprin-1 and Sun-1, impairs nuclear reorientation towards the synapse and leads to defects in actin organization at this level. Consequently, B cells are unable to internalize the BCR after antigen activation. Nesprin-1 and Sun-1-silenced B cells also fail to accumulate the tethering factor Exo70 at the center of the synaptic membrane and display defective lysosome positioning, impairing efficient antigen extraction at the immune synapse. Thus, changes in nuclear morphology and positioning emerge as critical regulatory steps to coordinate B cell activation.

## Introduction

Efficient uptake and processing of foreign antigens by B cells is critical for their complete activation (Carrasco & Batista, 2007; Junt et al, 2007). In lymph nodes, B cell activation is initiated by the engagement of the B cell receptor (BCR) with antigens displayed at the surface of neighboring cells, which triggers tyrosine kinase-dependent signaling cascades (del Valle Batalla et al, 2018; Seda & Mraz, 2015; Obino et al, 2017; Yuseff et al, 2013). BCR signaling is coupled to a rapid actin-dependent membrane spreading response, followed by an acto-myosin-dependent contraction phase, enabling B cells to concentrate antigens at the center of the immune synapse (Harwood & Batista, 2011; Fleire et al, 2006; Schnyder et al, 2011; Natkanski et al, 2013). Concomitantly, B cells re-position the MTOC to the center of the synaptic membrane, which acts as a landmark to guide the polarized recruitment of lysosomes, which upon secretion can facilitate antigen extraction (Vascotto et al, 2007; Yuseff et al, 2011a; Reversat et al, 2015). How B lymphocytes manage to position the MTOC and target lysosomes to precise domains of the immune synapse remains incompletely understood. Recently, the tethering factor, Exo70, a subunit of the exocyst complex, was shown to be involved in lysosome docking at the synaptic membrane in B cells. Exo70 is associated to the MTOC in resting B cells and becomes repositioned to the synaptic membrane upon activation, helping to promote lysosome tethering and fusion (Sáez et al, 2019).

B cells possess reduced cytoplasmic space, occupied mainly by their large nucleus (Lebien & Tedder, 2008), which is closely associated to the MTOC, from which it becomes uncoupled during activation (Obino et al, 2016). We thus sought to determine the impact of nuclear morphology in cell reorganization during immune synapse formation of B cells. Nuclear morphology and positioning have been shown to regulate diverse cellular functions, including signaling, gene expression (Duong et al, 2014), DNA repair or genome distribution (Zink et al, 2004; Roux & Burke, 2007; de las Heras & Schirmer, 2014), as well as cell shape (Chen et al, 2015) and migration (Lämmermann et al, 2008; Beadle et al, 2008; Calero-Cuenca et al, 2021). Nuclear size, form and positioning rely on the Linker of Nucleoskeleton and Cytoskeleton (LINC) complex (Hao & Starr, 2019; Calero-Cuenca et al, 2018; Chang et al, 2015). This complex is formed by two families of transmembrane proteins: KASH/Syne/Nesprin proteins, anchored to the outer nuclear membrane, where they are connected to cytoskeletal components; and Sun domain proteins inserted at the inner nuclear membrane, which are associated to lamins and chromatin (Crisp et al, 2006). Interactions between nuclear envelope proteins and the surrounding cytoskeleton regulate nuclear positioning and cell polarity (Chang et al, 2015). For instance, Nesprin-Sun complexes directly connect actin filaments and microtubules with the nucleus, to control both nuclear and MTOC repositioning during cell migration (Gomes et al, 2005). In activated T lymphocytes, Lamin A interacts with the Nesprin-Sun complex to promote MTOC repositioning towards the immune synapse (Rocha-Perugini & González-Granado, 2014).

Whether B lymphocytes adjust their nuclear morphology to promote immune synapse organization, has not been addressed. In this work, we reveal that, upon activation with immobilized antigens, B cells re-orientate their nuclear groove, in an actin and microtubule dependent manner, towards the antigen contact site, where the immune synapse is formed. Silencing the expression of nuclear envelope proteins, Nesprin-1 or Sun-1, decreased cytoskeleton-nuclear connections and impaired nuclear reorientation to the immune synapse. Strikingly, these cells displayed an extremely disorganized immune synapse, characterized by decreased actin cytoskeleton levels and deficient internalization of the BCR at the center of the synapse. Additionally, Nesprin-1 and Sun-1 silenced B cells mislocated the tethering factor Exo70 and were unable to concentrate lysosomes to the center of the synaptic interface. As a consequence of the disorganized immune synapse B cells decreased their antigen extraction capacity. Thus, our results highlight how B cells adjust nuclear morphology to cytoskeleton rearrangements at the immune synapse to orchestrate lysosome recruitment and enable them to acquire their antigen extraction and processing functions.

## Results

### B cells change their nuclear morphology and re-orientate their nuclear groove toward the immune synapse

We first evaluated whether B cells change their nuclear morphology upon activation with immobilized antigens. To this end, we labeled the nuclear lamina of B cells under resting or activating conditions, using anti-Lamin-B, which detects the only Lamin expressed in these cells (Jansen et al, 1997). We also labeled microtubules (α-tubulin) and actin (phalloidin) and performed 2D and 3D reconstructions (Fig. EV1A). Our analysis revealed that in resting cells the nucleus occupied 70% of the total cell volume (Fig. EVB,C). Resting B cells, displayed a lobular shaped nucleus (between 3 or 4 lobes), where the MTOC was closely associated inside the main nuclear groove or within a principal intra-lobular space, similar to recently described studies in hematopoietic cells (Biedzinski et al, 2020)(Fig. EV1A-B). We next evaluated nuclear morphology under activating conditions by incubating B cells with 3μm beads coated with antigen, which mimics the formation of an immune synapse (Yuseff et al, 2011a). During early activation times (30-60 min), B cells re-positioned their nuclear groove towards the antigen (Fig. 1A-C) which was coupled to the polarization of the MTOC (Fig. EV1D,E). At later time points, we found that the antigen-coated bead became accommodated within the main nuclear groove near the MTOC, suggesting that B cells adapt the position and morphology of the nucleus to create a space to bring the MTOC closer the synaptic interface. Changes in the area of the main nuclear groove were also observed in activated B cells, which decreased after 30 min of activation and were restored to the original size after 60 min (Fig. 1D). Interestingly, we observed that B cells interacting with multiple antigen-coated beads displayed larger nuclear grooves, suggesting that they adjust this area to the size to the surface containing the immobilized antigen (Fig. 1E,F). To further characterize nuclear reorientation, we used another experimental setup where the antigen was fixed to coverslips, thereby allowing us to evaluate movements or rotation of the nucleus to the antigen contact site. B cells seeded onto antigen-coated dishes for different time points were labeled for lamin B/Hoechst and actin (phalloidin). Similarly to our observations with antigen coated beads, at early time point of activation, the nuclear groove area of B cells decreased (50%) and then increased restoring its size after 60 min of activation (Fig. EV1F) suggesting that reduction in the nuclear groove area occurs concomitantly with nuclear rotation. Noticeably, we observed that the nuclear groove became fully reoriented towards the central region of the immune synapse at 30 min (Fig. 1G,H), where lysosomes progressively accumulated (Fig. EV1G). These results highlight how B cells change their nuclear morphology during different stages of activation and how lysosomes are accommodated at the central region between the nucleus and the synaptic membrane inside the nuclear groove.

**Figure 1.**
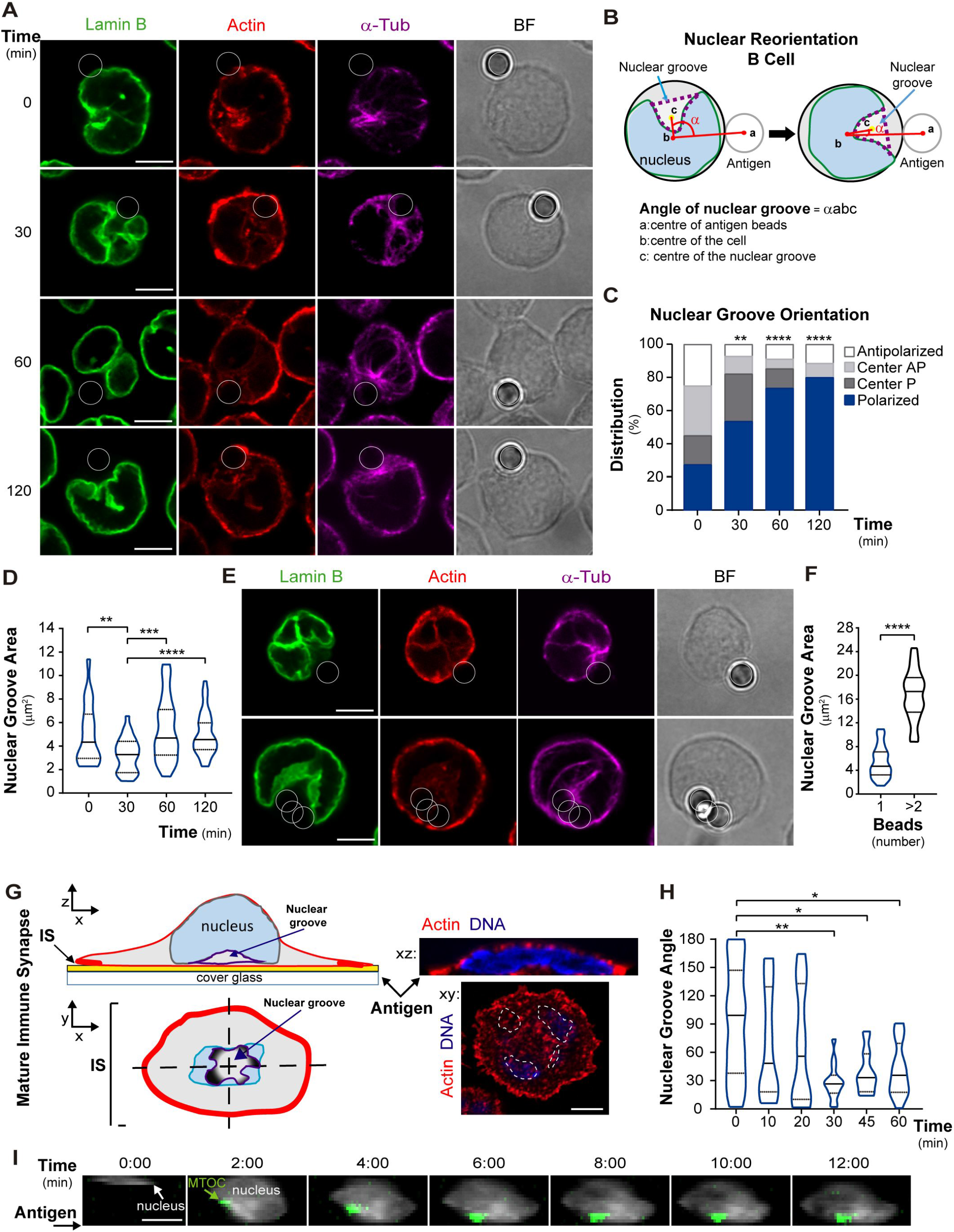
Nucleus is reoriented toward the immune synapse of B cells. A. Confocal representative images of B cells stained for the nucleus (lamin B in green), actin (phalloidin in red) and microtubules (α-tubulin in magenta). Cells were incubated with antigen coated beads for 0, 30, 60 and 120 min. White circles indicate bead position. Scale bar 5 µm. B. Scheme depicting the method used to quantify nucleus orientation. We manually delimited the space left by lamin B staining in the principal nuclear groove and the quantification was performed as indicated in Methods. C. Quantification of nuclear groove orientation measuring percentage of cells that have nucleus toward the antigen coated beads, as described in Methods; n≥90; three independent experiments. Kruskal-Wallis test with Dunnś multiple comparisons test was performed for statistical analyses. N≥60, *p<0.05, **p<0.01; ****p<0.0001. D. Quantification of nuclear groove area in 2D. B cells were activated as described in A. Statistical analyses, as C. N≥62, **p<0.01; ***p<0.001,****p<0.0001. E. –F. Fluorescence images of B-cells activated and stained for the nucleus and cytoskeleton as described in A, A Cell interacting with multiple activating beads, increasing antigen area, is shown. Scale bar 5 µm. Left, quantification of the nuclear groove area in these conditions. Unpaired test, ****p<0.0001. N=40. G. Scheme and images describing mature immune synapse in xz and xy, in B cells seeded on antigen coated dishes. Staining was nucleus (Hoechst in blue), actin (phalloidin in red). Nuclear groove is at the center of the immune synapse. White line indicates the position of the nucleus H. Quantification of nuclear groove orientation in B cells seeded on antigen coated dishes for 0, 10, 20, 30, 45 and 60 and 120 min. Statistical analyses, as C. N≥35, *p<0.05, **p<0.01. I. Visualization of the nucleus (Hoechst in blue) and MTOC (centrin-GFP) of B cells seeded on antigen coated dishes (filmed each 2 min). Scale bar 5 µm.

In order to monitor dynamic changes in nuclear morphology and MTOC polarization, simultaneously, during B cell activation, we next performed live imaging of centrin-GFP expressing B cells labeled with Hoechst, using the same experimental setup, and filmed every 2 min by epifluorescence microscopy. Our results show that as the MTOC became positioned towards the synaptic membrane, the nucleus rotates and changes from a circular form (before MTOC reorientation) to a hat shape (after MTOC reorientation) (Fig. 1I). Thus, engagement of B cells with immobilized antigens triggers changes in nuclear morphology and reorientation of their nuclear groove to the center of the immune synapse.

### Cytoskeleton-dependent nuclear reorientation to the Immune Synapse of B cells

Nuclear morphology is known to be controlled by both actin and microtubules (Gundersen & Worman, 2013; Wilson & Holzbaur, 2012; Lele et al, 2018; Cadot et al, 2012; Roman et al, 2017; Starr & Fridolfsson, 2010), thus we verified their association to the nucleus of B cells. To characterize cytoskeleton-nucleus connections, resting or activated B cells were labeled with Lamin B, actin (phalloidin) and microtubules (α-tubulin). We observed that under resting conditions, actin cytoskeleton and microtubules were strongly associated to the nucleus and inside the main nuclear groove, nascent microtubules emerging from the MTOC closely surrounded each nuclear lobe (Fig. EV2A). We could also visualize the actin pool surrounding the MTOC, which was enriched inside the nuclear groove. Interestingly, actin and microtubules were associated with nuclear groove borders and were enriched in lobe vertexes and curvatures and also on top of the nucleus which could regulate nuclear positioning. Orthogonal views of activated B cells, revealed increased associations between actin and the nucleus (Fig. 2A), suggesting that nucleus-cytoskeleton connections are modified upon antigen engagement. Importantly, studies in primary B cells, whose cellular and nuclear diameters are 6.5 μm, and 5.36 μm, respectively (Fig. EV2B,C), displayed smaller nuclear grooves (0.4 μm^2^ area) (Fig. EV2D), and behaved similarly to the B cell line, in terms of nuclear re-orientation (Fig. EV2E).

**Figure 2.**
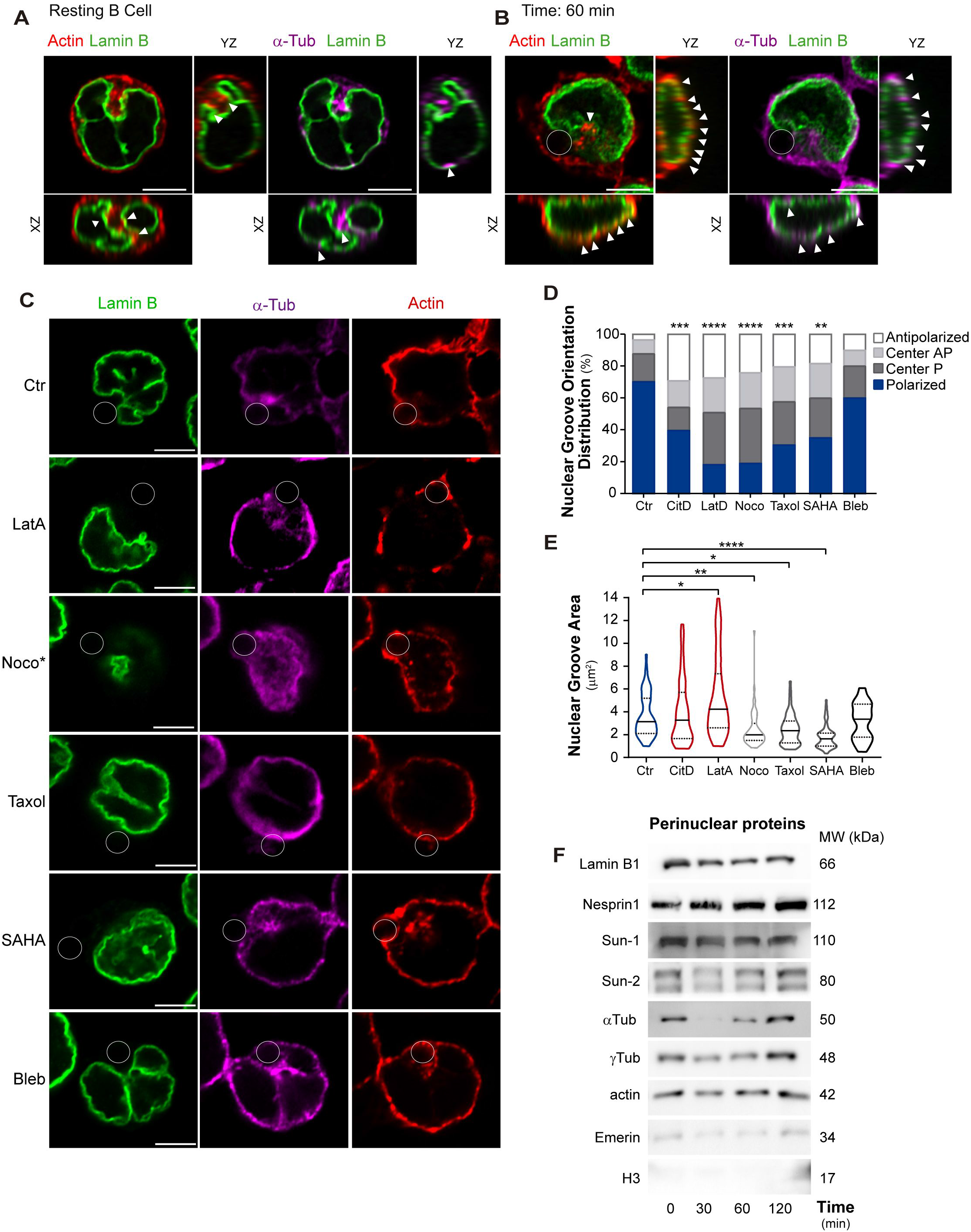
Cytoskeleton stability impacts nuclear shape to the Immune Synapse of B cells. A. Confocal images showing cytoskeleton actin (red) and microtubules (magenta) highly associated to nuclear morphology (lamin B, green) in resting in B cells. Arrowheads indicate cytoskeleton-nucleus connections. B. Representative confocal images of B cells incubated with antigen coated beads for 60 min. Cells were stained as in A. C. –E Confocal images showing the immune synapse plane of B cells incubated 60 min with antigen coated beads. After 10 min of activation, cells were treated with nocodazole, taxol, latrunculin A, cytochalasin D, SAHA or Blebbistatin. *Noco, immune synapse plane was far from the rest of the B cell. White circles indicate bead position. Quantification of the percentage of cells that have orientated the nucleus was as indicated in Methods. Quantification of nuclear groove area in 2D. Kruskal-Wallis test with Dunnś multiple comparisons test was performed for statistical analyses; n≥60 cells from two independent experiments, *p<0.05,**p<0.01,***p<0.001, ****p<0.0001. F. Western blot showing protein levels in perinuclear fractions purified from B cells activated at different time points. The perinuclear fractions contain: Nesprin-1, Emerin, Sun-1, Sun-2, αTub, γTub, actin, but no histone-3 (H3). Representative of 3 independent experiments.

We further assessed the role of the cytoskeleton in nuclear morphology by evaluating the effects of drugs that perturb microtubules (Nocodazole, Taxol) or actin (Cytochalasin D and Latrunculin A). Nuclear morphology was evaluated in resting B cells preincubated with each drug for 30 min and labeled for Lamin B. 3D reconstructions of these cells revealed that all drugs that perturbed either microtubules or the actin cytoskeleton, decreased the total nuclear volume (Fig. EV3A), highlighting how nuclear morphology in B cells is tightly coupled to cytoskeleton networks, as observed in other cell types. We next performed these studies in activating conditions, where cytoskeleton perturbing drugs as well as SAHA (increases microtubules acetylation) and Blebbistatin (myosin II inhibitor), were added 10 min after activation. Our results show that the orientation of the nuclear groove towards the immune synapse was significantly impaired when intervening with either microtubules or actin cytoskeleton (Fig. 2C,D and Fig. EV3B). Interestingly, the size of the nucleus also changed and increased when actin depolymerization was induced by LatA, whereas it decreased in the presence of drugs that disrupted or fixed microtubules (Fig. 2F). These results were correlated to those obtained for the MTOC polarization analysis (Fig EV3C) confirming the high association with the nuclear groove morphology and MTOC reorientation. No effects on nuclear morphology or reorientation were observed upon treatment with Blebbistatin, indicating that they did not rely on myosin II activity.

Considering that actin and microtubules regulate nuclear morphology and orientation in B cells, we next asked whether such interactions were regulated during activation. This was evaluated by assessing changes in the composition of nucleus-associated proteins in resting and activated B cells. To this end, nuclear, perinuclear and cytoplasmatic fractions were isolated by subcellular fractionation of lysates obtained from resting B cells (Fig. EV3D) and under different activation times (Fig. EV3E). We focused on the perinuclear fraction, which contains proteins bound directly or indirectly to the inner or the outer nuclear membrane (Shaiken & Opekun, 2014). Our results show that total perinuclear protein levels increased during the time course of activation (Table 1), reaching about 30 times more after two hours of activation with respect to non-activated cells (from 0.16% to 2.99%), suggesting that upon activation, B cells redistribute proteins to the vicinity of the nucleus. Analysis of cytoskeletal components in perinuclear fractions (Fig. 2F) revealed stable actin levels and only a slight decrease after 30 min of the activation, which were later reestablished. This result is in agreement with a previous report showing that B cells deplete actin from the perinuclear region in order to promote MTOC repositioning to the immune synapse (Ibañez-Vega et al, 2019; Obino et al, 2016). In contrast, levels of Lamin B in the perinuclear fraction gradually decreased during activation, suggesting that Lamin B became less coupled to the nuclear membrane. Perinuclear levels of α-Tubulin strongly decreased upon activation as well as γ-Tubulin levels, which diminished after 30 and 60 min of activation and were reestablished after 120 min. These results suggest that, in resting B cells, microtubules are associated to the nucleus and become less coupled from this organelle upon activation when this is rotating. The higher levels of γ-Tubulin detected in perinuclear fractions after longer time points of activation, most likely reflect the tight association of the MTOC with the nuclear membrane when the nucleus is fully reoriented to the immune synapse.

**Table 1:** Subcellular distribution of total proteins during B cell activation. B cells were activated for indicated times point and the percentage of proteins contained in the cytoplasm, nucleus and perinucleus obtained by subcellular fractionation was calculated (as described in Methods).

| % of Total<br>Lysate | Fractions |  |  |
| --- | --- | --- | --- |
|  | Cytoplasm | Perinucleus | Nucleus |
| t = 0 min | 82.85 | 0.16 | 16.98 |
| t = 30 min | 76.63 | 0.10 | 27.28 |
| t = 60 min | 80.56 | 1.52 | 16.71 |
| t = 120 min | 77.35 | 2.99 | 21.10 |

Additionally, we found that the levels of nuclear membrane associated proteins, Nesprin-1, Sun-1, Sun2 (Starr & Fridolfsson, 2010) and emerin (Chang et al, 2013; Storey et al, 2020), in perinuclear fractions did not change over the time course of activation, suggesting that their associations to the nucleus are stable as opposed to actin, microtubules and lamin B, which presented dynamic nuclear association-disassociations. Overall, our results show that B cells regulate the association of actin and microtubules to the nuclear envelope, which promote nuclear reorientation toward the immune synapse.

### B cells depend on conserved nuclear-cytoskeleton proteins, Nesprin-1 and Sun-1, to regulate nuclear morphology

Nesprin and Sun proteins regulate nuclear shape, size and positioning in different cell types and frequently form complexes in order to perform these effector functions (Duong et al, 2014; Lombardi et al, 2011; Méjat & Misteli, 2010). We thus evaluated whether such complexes could be formed during B cell activation. For this purpose, we performed immunoprecipitation assays with nuclear membrane proteins, which revealed that under resting conditions, Nesprin-1 formed a complex with Sun-1 but not with Sun-2, and upon activation, Nesprin-1 and Sun-1 interactions increased during a similar time frame where Nesprin-1 becomes more associated to actin (Fig. 3A). These results indicate that complexes formed by nuclear envelope proteins, Nesprin-1 and Sun-1 interact with actin and could regulate nuclear morphology in B cells. To evaluate their role in regulating nuclear morphology during B cell activation, we silenced Nesprin-1 or Sun-1, achieving a 60% decrease in their expression levels (Fig. EV4A) and performed immunofluorescence staining of the nucleus (lamin B) and actin. Under resting conditions, B cells silenced for Nesprin-1 or Sun-1 displayed an actin cortex which was more dispersed and separated from the nucleus, compared to control cells (Fig. 3B), suggesting that actin-nuclear membrane connections were lost. To confirm this, we performed 3D analysis of confocal images of B cells stained for the nucleus and actin, using SCIAN-Soft/IDL software. Our analysis revealed that the cortical actin cytoskeleton of control cells formed a sphere that covered the nucleus, while perinuclear actin was concentrated within the nuclear groove (Fig. 3C). In contrast, Nesprin-1- or Sun-1-silenced B cells lost the spherical form of the cortical actin cytoskeleton and displayed ruffles which extended away from the nucleus, and decreased levels of perinuclear actin within the nuclear groove. We next evaluated whether the loss of actin within this region had an impact in the nuclear groove area. Indeed, we observed that under resting conditions, Nesprin-1 silenced B cells increased their nuclear groove area (Fig. 3D and EV4B) which was also observed at early time points of activation compared to control counterparts, showing no significant changes after 60-120 min of activation (Fig. EV4B). On the other hand, Sun-1-silenced B cells did not display major differences in their nuclear groove area compared to control cells, which was only significantly reduced at later times of activation (Fig. 3D and EV4B). Additionally, we evaluated whether Nesprin-1 and Sun-1 regulate nuclear rotation. B cells were seeded onto antigen-coated dishes and we quantified the angle of the two lobules surrounding the nuclear groove facing the immune synapse. This analysis revealed that Sun-1 and Nesprin-1 silenced B cells exhibited impaired nuclear reorientation (Fig. 3E,F).

**Figure 3.**
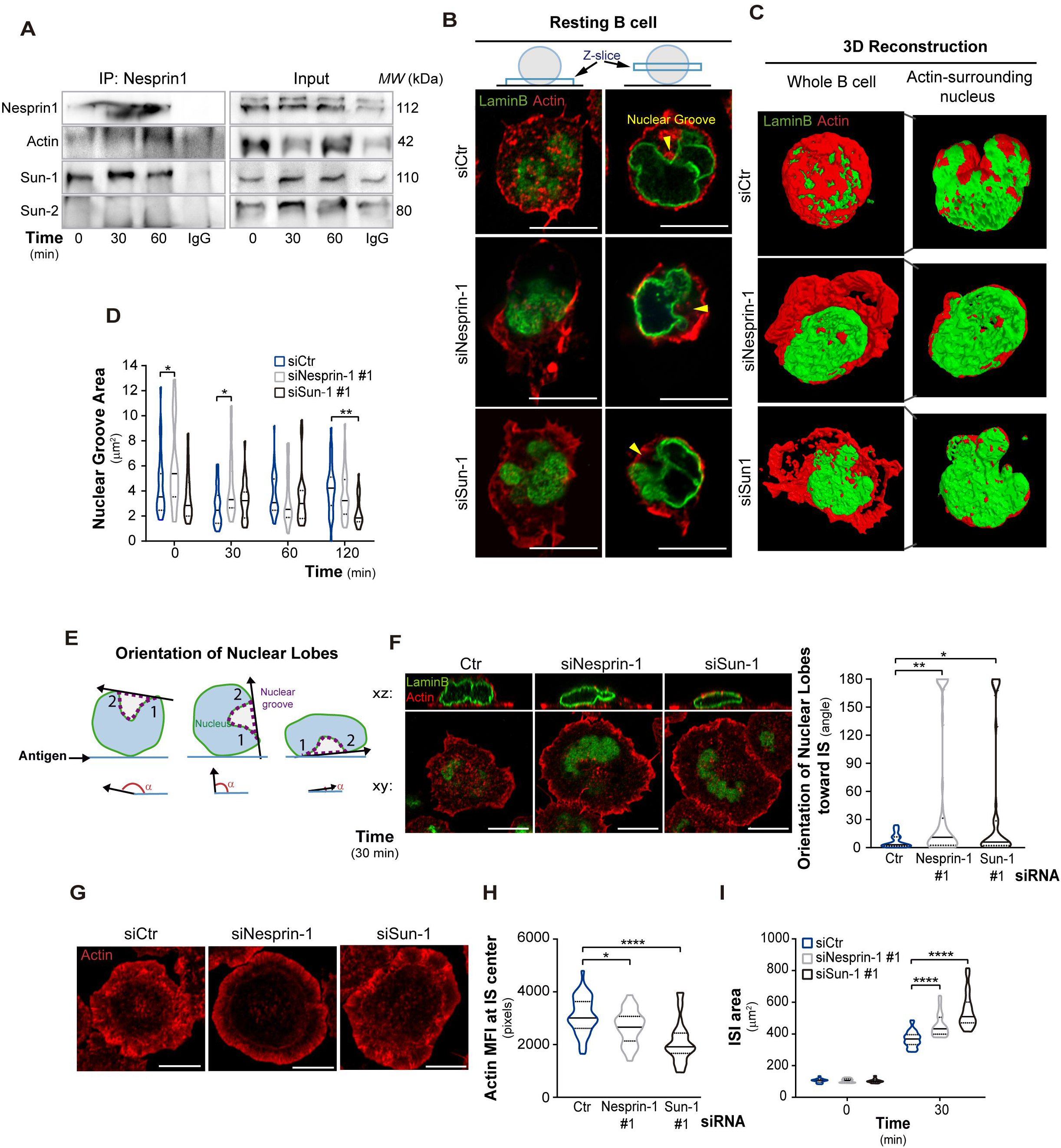
Nesprin-1 and Sun-1 regulate nuclear shape during B cell activation. A. Nesprin-1 immunoprecipitation (IP) assay to detect the formation of LINC complexes (Nesprin-Sun) in resting or activated B cells for indicated times. Representative of 2 independent experiments. B. Representative confocal images of Control, Nesprin-1 and Sun-1 -deficient B cells, which show 2 different planes: left, glass cover plane; and right, nuclear groove plane. Nucleus (lamin green) and actin (phalloidin red) demonstrate detachment of actin cortex from the nucleus. Yellow arrowheads indicate nuclear groove. Scale bar: 10 μm C. 3D reconstructions images of the nucleus staining with lamin B (green) and actin (red). Left image shows complete actin and lamin B signals; and right, logical filter is applied on actin signal showing only actin co-localizes with lamin B. D. Quantification of nuclear groove area of Control or Nesprin-1 and Sun-1 -silenced B cells incubated with antigen coated beads for indicated times. Kruskal-Wallis test with Dunnś test was performed for statistical analyses; n≥40 cells from two independent experiments, *p<0.05, **p<0.01. E. –F Quantification of nuclear groove rotation of Control or Nesprin-1 and Sun-1 -silenced B cells incubated on antigen coated dishes for 30 min. Scheme depicting the method used to determine the angle of two nuclear lobes that surround the nuclear groove. Representative confocal images showing the immune synapse plane and Resliced confocal image to determine nuclear lobes angles in 2D. Quantification of nuclear Lobes orientation. Two-way ANOVA with Sidak’s multiple comparison test; n≥50 cells from two independent experiments **p<0.001, *p<0.05. G. –I. Representative confocal images showing the immune synapse plane of Control or Nesprin-1 and Sun-1 -silenced B cells seeded onto antigen coated dishes, for 30 min. Actin (red). Quantification of actin at the center of the immune synapse. Statistical analyses, as D; n≥45 cells from three independent experiments *p<0.05, ****p<0.0001. Quantification of the immune synapse spreading area. Statistical analyses, as D; n≥60 cells, ****p<0.0001.

Considering that Nesprin-1 and Sun-1 are required to maintain actin connected to the nuclear groove in resting cells, we asked whether they could also regulate actin reorganization at the immune synapse, where the nucleus becomes tightly re-positioned. To this end, we quantified actin at the center of the immune synapse where the nuclear groove sits and measured peripheral actin, which limits the area of the synaptic membrane. Our results show that Nesprin-1 and Sun-1-silenced B cells display significant defects in actin reorganization at the immune synapse. Such defects include decreased actin levels at the center of the synapse (Fig. 3G,H) and the formation of large peripheral lamellipodia. Consequently these cells also displayed increased spreading areas, 100 and 150 μm2 compared to control cells (Fig. 3G,I). Altogether these results show that Sun-1 and Nesprin-1 regulate actin organization at the nuclear interface as well as at the immune synapse.

### Nesprin-1 and Sun-1 are required for efficient antigen extraction but not for MTOC or lysosome polarization

Having shown that Nesprin-1 and Sun-1 connect actin to the nuclear groove, we next assessed their role in regulating the polarization of key organelles, such as the MTOC and lysosomes, which rely on changes in actin dynamics at the nuclear interface. MTOC and lysosomes were labeled in resting and activated B cells silenced for Nesprin-1 or Sun-1, and their repositioning towards the antigen coated-beads was evaluated. Our results show that Nesprin-1- and Sun-1-silenced B cells did not display significant defects in MTOC (Fig. EV5A,B,C) or lysosome polarization (Fig. EV4B,C). However, we observed defects in their antigen extraction capacity (Fig. 4A), which was evaluated by measuring the amount of antigen (OVA) remaining on beads in contact with activated B cells. Indeed, Nesprin-1- and Sun-1-silenced B cells extracted less antigen compared to control cells (Fig. 4B,C). To reveal the cause of defective antigen extraction, we analyzed the distribution of lysosomes at the synaptic interface of Nesprin-1- and Sun-1-silenced B cells in more detail. Upon activation, we observed that these cells were unable to form lysosome LAMP1+ rings that typically surround the activating beads, suggesting that lysosome docking and secretion at the synaptic interface was impaired, thereby explaining their lower antigen extraction capacity.

**Figure 4:**
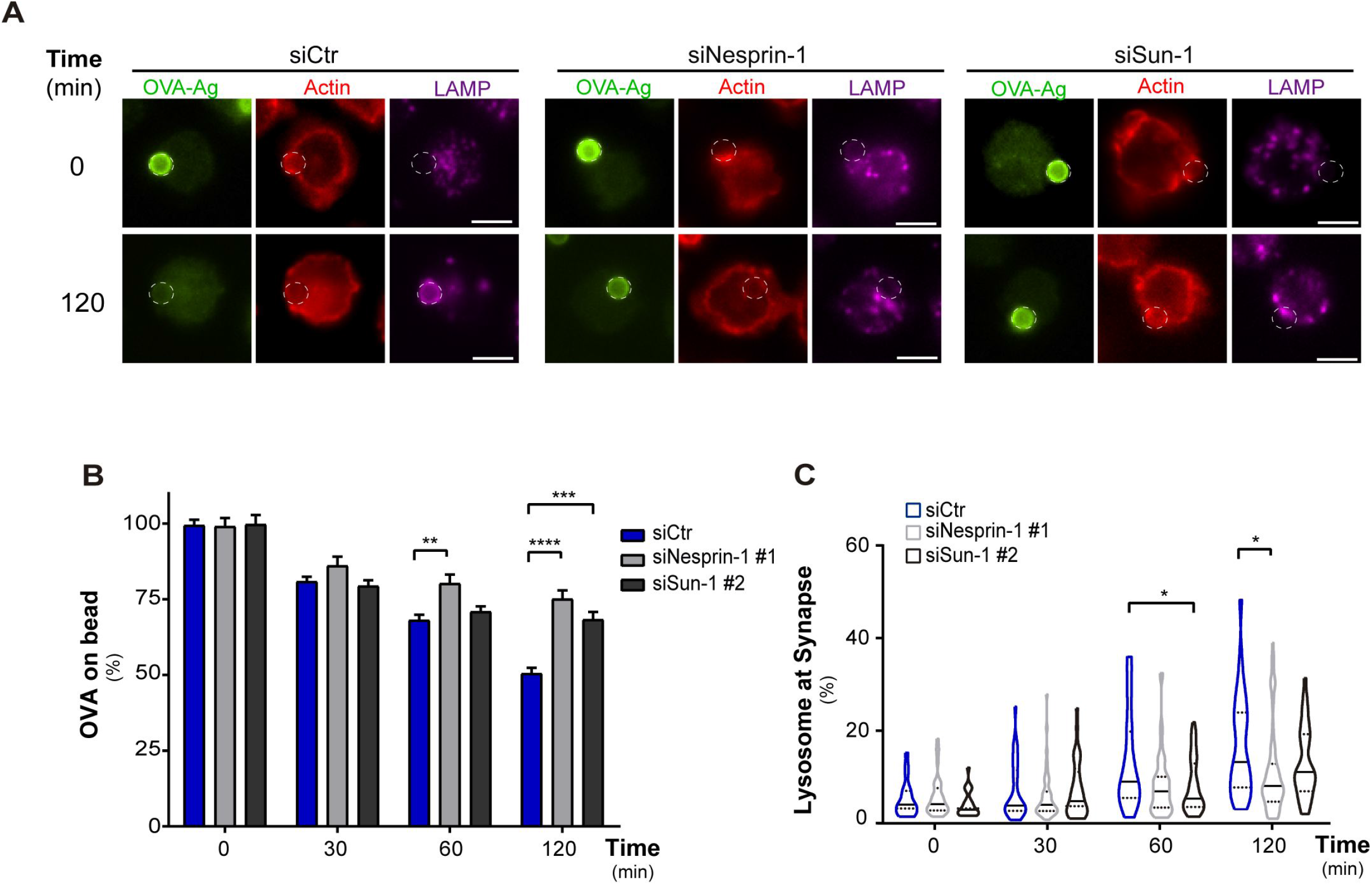
Antigen extraction relies on Nesprin-1 and Sun-1. A. Epifluorescence images of Control and Nesprin-1 and Sun-1 silenced B cells incubated with antigen coated beads for 0 or 120 min. Cells were stained against OVA (Green), actin (red) and LAMP1 (magenta). White circles indicate bead position. Scale bar 5 μm. B. Quantification of percentage of OVA remaining on antigen coated beads. Kruskal-Wallis test with Dunn’s multiple comparisons test was performed for statistical analyses. n=80, **p=0.001;***p=0.0006; ****p<0.0001. Mean with SEM line are shown. C. Quantification of LAMP1+ rings surrounding antigen coated beads. Statistical analyses, as B. n=80, *p=0.05.

### Nesprin-1 and Sun-1 are required for B cells to establish an organized immune synapse

Given that Sun-1/Nesprin-1-silenced B cells displayed defects in lysosome accumulation around the antigen-coated beads, as well as in actin organization at the synaptic interface, we next asked whether the overall organization of the immune synapse was altered under these conditions. To this end, control or Sun-1/Nesprin-1-silenced B cells were activated on antigen-coated dishes and the distribution (x y z) of BCR, microtubules/MTOC, lysosomes and the nucleus were evaluated at the immune synapse. Our results show that the BCR of Nesprin-1- and Sun-1-silenced B cells remained at the center of the synaptic membrane and was not internalized as observed in control cells (Fig. 5A,B and EV6A (z)). The MTOC of these cells can still polarize but its signal was increased at the synaptic membrane compared to control cells (Fig. 5C,D). Although, the lysosomes of Nesprin-1- and Sun-1-silenced B cells were polarized (z) to a similar extent compared to control cells (Fig. EV6B), these were completely mispositioned respect to the synapse center (x y) (Fig. 5E,F). Notably, the nucleus of these cells was also displaced with respect to the synapse center and closer to the synaptic membrane (Fig. 5E,G), suggesting that nuclear and lysosome positioning are tightly coupled. Thus, silencing nuclear envelope proteins alters the distribution of BCR and MTOC localization and also lysosome recruitment towards the center of the immune synapse, highlighting a functional link between the regulation of nuclear morphology and immune synapse organization.

**Figure 5:**
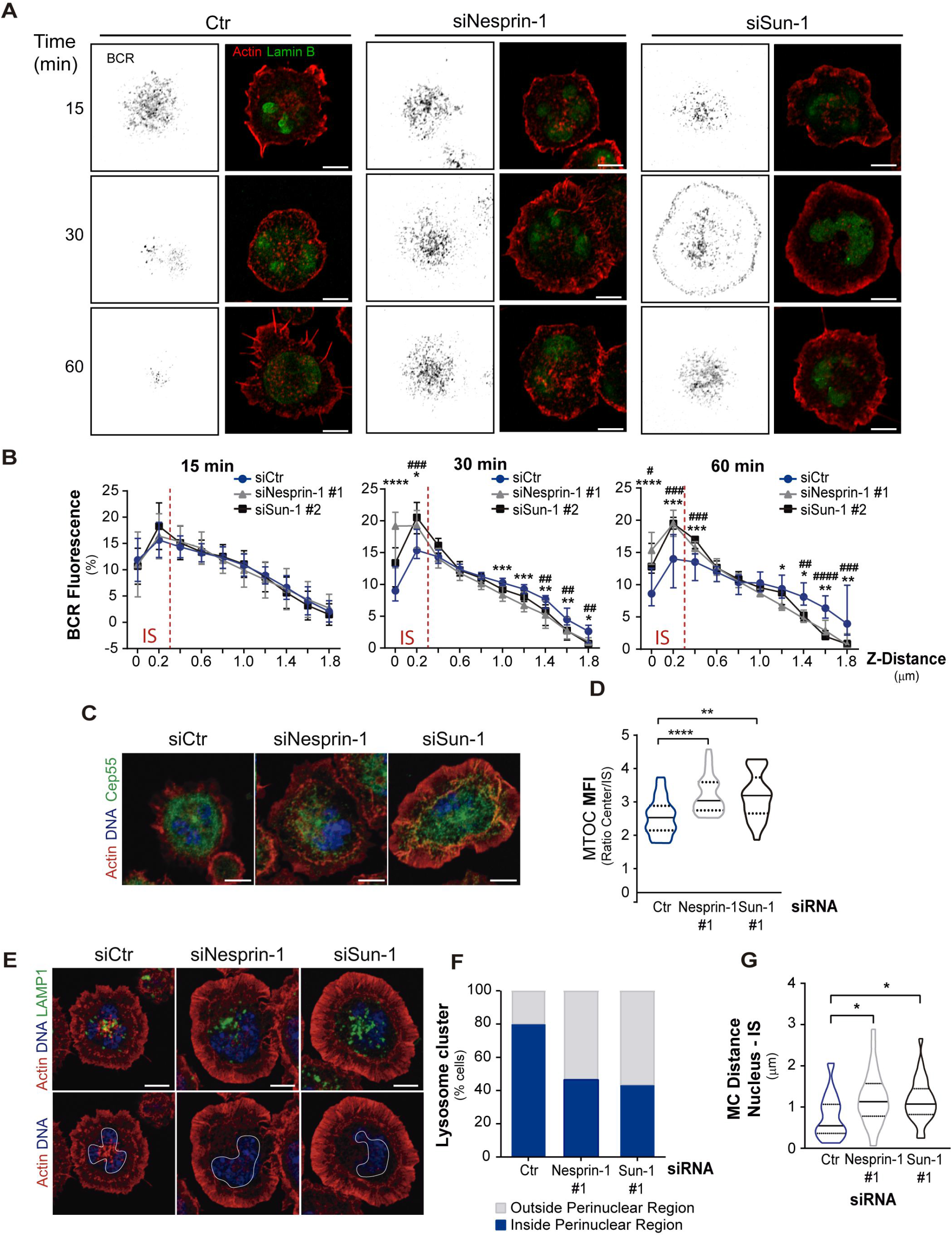
Nesprin-1 and Sun-1 regulate Immune synapse organization. A. Confocal images of Control and Nesprin-1 and Sun-1 silenced B cells activated on antigen coated dishes for 15, 30 and 60 min. BCR (grey color), actin (red) lamin B (green) Scale bar 5 µm. B. Quantification of BCR fluorescence intensity at immune synapse (0-0.2 μm) and intracellular localization (0.4-2 μm) following z distance. Mixed-effects analysis and Dunnettś multiple comparisons test. n>30. * and # are used to show statistical differences between control and Nesprin-1 or Sun-1 silenced cells, respectively. */# p<0.05, **/##p<0.01, ***/###p<0.001,****/####p<0.0001. Mean with 95% CI line are shown. C. -D. Confocal images of Control and Nesprin-1 and Sun-1 silenced B cells seeded on antigen coated dishes for 30 min. Microtubules/MTOC (cep55 green), actin (red) and nucleus (blue). Scale bar 5 µm. Quantification of MTOC at the immune synapse. Kruskal-Wallis test with Dunn ’s multiple comparisons test was performed for statistical analyses. n>52,**p<0.01,***p<0.001. E. -G. Confocal images of Control and Nesprin-1 and Sun-1 silenced B cells seeded on antigen coated dishes for 30 min, upper panel show actin (red) and nucleus (DNA, blue); lower panel, plus lysosomes (LAMP1, green). White lines indicate the position of the nucleus. Scale bar 5 µm. Quantification of percentage of cells that have the principal cluster of lysosomes inside or outside of a perinuclear region at the immune synapse. n>60. Quantification of nuclear dislocation from the immune synapse (distance between nuclear center and immune synapse center) Statistical analyses, as D. n>60, *p<0.05.

### Exo70 is necessary to maintain nuclear positioning at the immune synapse and its localization rely on Nesprin-1 and Sun-1 in B cells

To obtain mechanistic insights underlying the altered positioning of lysosomes at the immune synapse in Nesprin-1 or Sun-1-silenced B cells, we focused on Exo70, a subunit of the exocyst complex described in trafficking and organelle biogenesis in other cell types, it was also involved in the tethering of secretory vesicles to specific domains of the plasma membrane in activated B cells (Hertzog et al, 2012; Sáez et al, 2019; Zhu et al, 2019). To evaluate whether Nesprin-1 or Sun-1-silenced B cells had defects in mobilizing Exo70 to the synaptic membrane, we evaluated its localization in resting and activating conditions in these cells. Using imaging analysis, we observed that Nesprin-1 and Sun-1 silenced B cells had lower levels of Exo70 associated to the MTOC under resting conditions (Fig. 6A,B) Upon activation Exo70 was distributed in a more dispersed fashion throughout the synaptic membrane, compared to control cells (Fig. 6C,D). These results suggest that Nesprin-1 and Sun-1 regulate the association of Exo70 to the MTOC and its further localization to the center of the immune synapse. Thus, the altered distribution of Exo70 in Nesprin-1 and Sun-1 silenced cells could explain why lysosomes fail to accumulate at the center of the synapse under these conditions.

**Figure 6:**
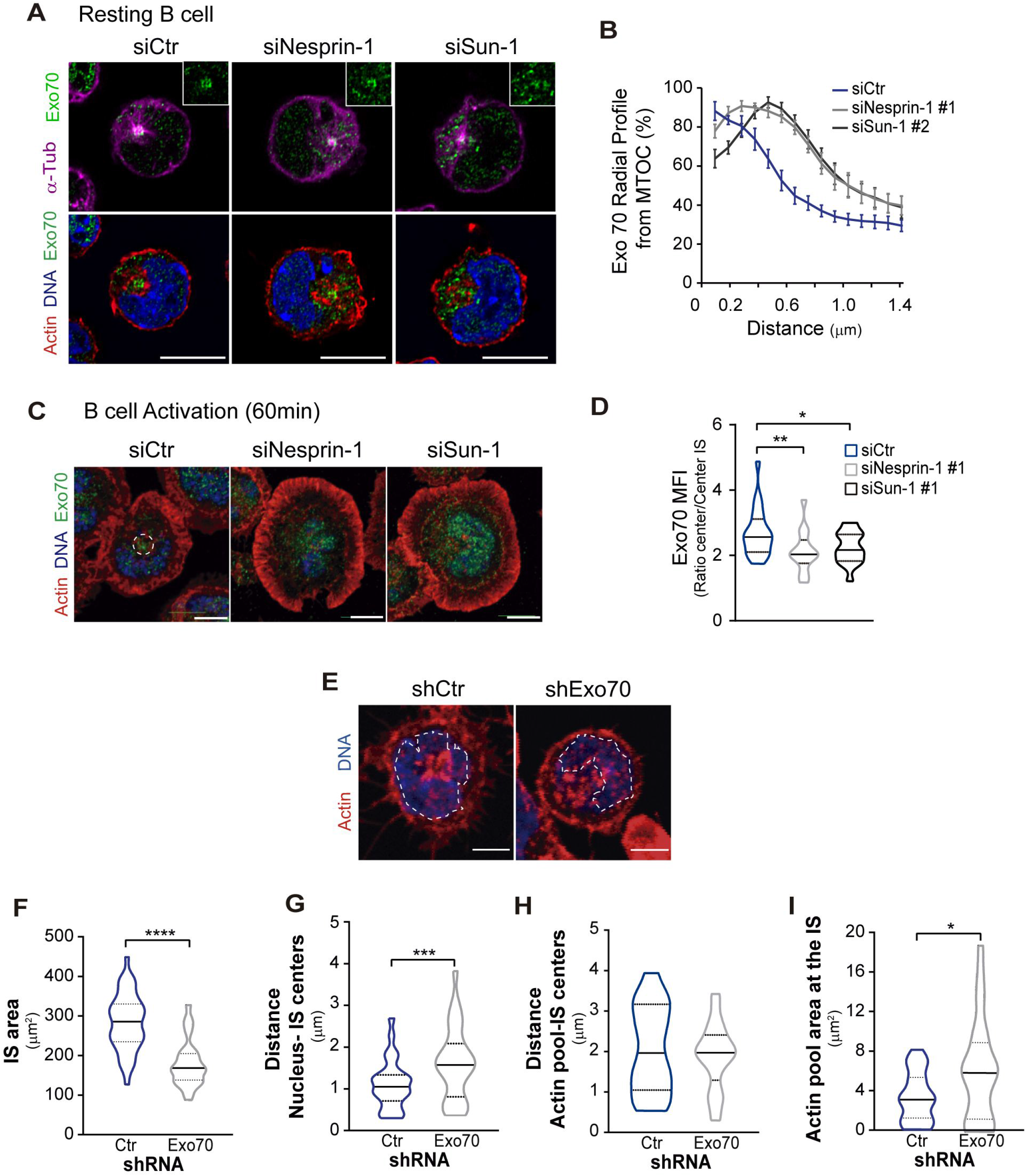
Exo70 localization is controlled by Nesprin-1 and Sun-1 in B cells. A. -B. Confocal images of Control and Nesprin-1 and Sun-1 silenced B cells in resting conditions, showing Exo70 (green) with: upper panel, microtubules/MTOC (α-Tub, magenta); and lower panel, actin (red) and nucleus (DNA, blue). Quantification of the Exo70 distribution (radial profile) from MTOC. Mixed-effects analysis and Dunnettś multiple comparisons test. n>30. * and # are used to show statistical differences between control and Nesprin-1 or Sun-1 silenced cells, respectively. */# p<0.05, **/##p<0.01, ***/###p<0.001, ****/####p<0.0001. Mean with 95% CI line are shown. C. -D. Confocal images of Control and Nesprin-1 and Sun-1 silenced B cells activated on antigen coated dishes for 30 min. Exo70 (green), actin (red) and DNA (blue). Quantification of the ratio of Exo70 in the central region vs the periphery at the immune synapse. Kruskal-Wallis test with Dunnś multiple comparisons test was performed for statistical analyses. n=40, *p<0.05. E. - I. Confocal images of Control and Exo70 silenced B cells activated on antigen coated dishes for 60 min. Actin (red) and nucleus (blue). White lines indicate the position of the nucleus. Quantification of the area of the immune synapse. Mann-Whitney test, ****p<0.0001; n=50. Quantification of nuclear dislocation from the immune synapse (distance between nuclear center and immune synapse center). Unpaired t test, ***p=0.0003; n=50. Quantification of the actin pool with respect to the center of the immune synapse. Statistical analyses, as F; n=50 no significantly different. Quantification of actin pool area at the immune synapse. Statistical analyses, as F, *p=0.0485; n=40.

**Figure 7:**
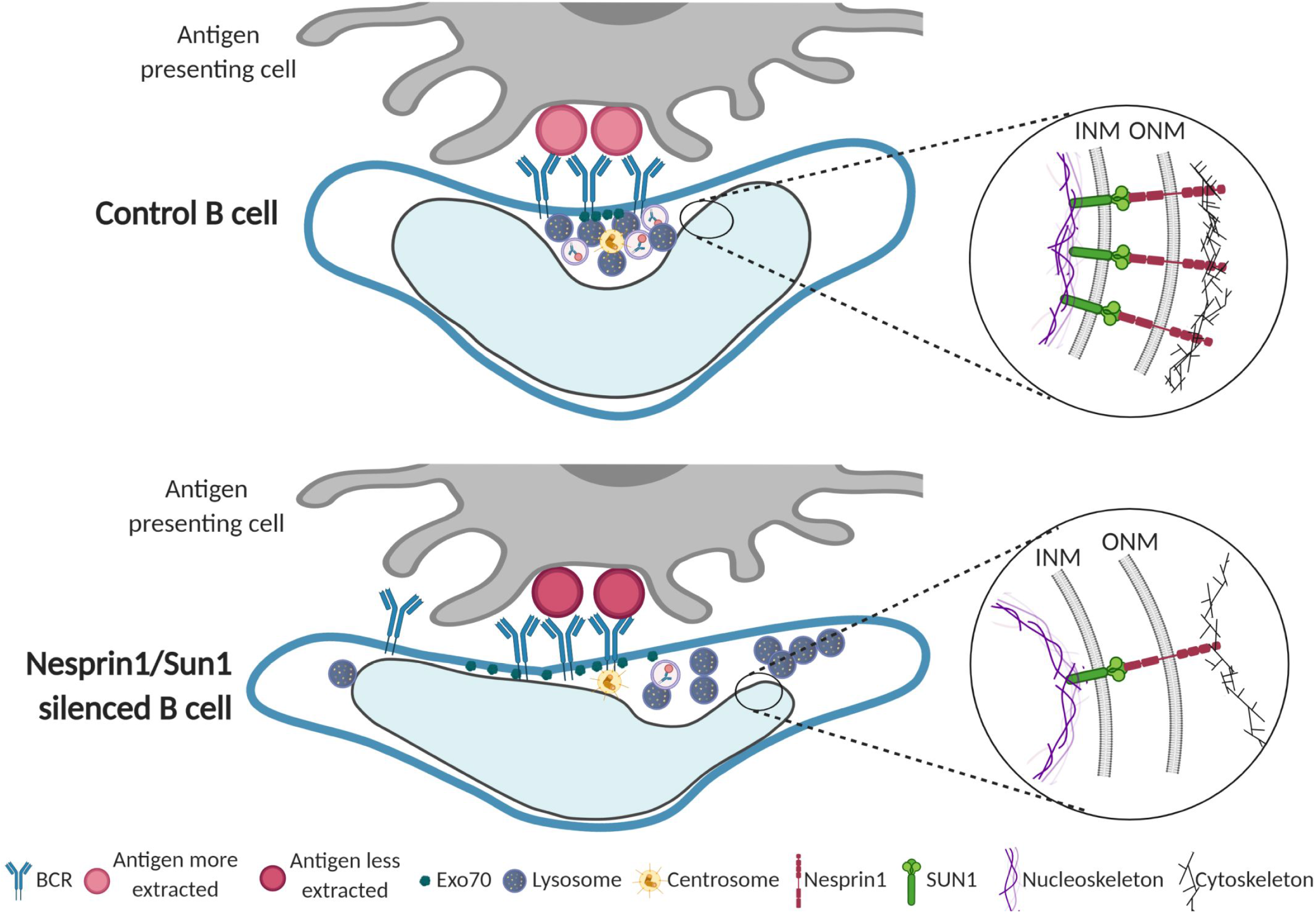
Model of the Immune synapse organization controlled by the nuclear envelope proteins Nesprin-1 and Sun 1. Upon activation, the immune synapse formed by the B cell and the antigen presenting cell is well organized. In the center of the immune synapse, the MTOC, Exo70 and the lysosome are recruited and antigen-BCR complexes are clustered and internalized. This region is formed by the positioning of the nuclear groove, which orchestrates cytoskeleton remodeling and membrane trafficking. In Nesprin-1 and Sun-1 silenced B cells, the nuclear groove is not well reoriented because actin nucleus connection are lost, disorganizing the immune synapse impaired BCR clustering at the center of the immune synapse. Exo70 fails to concentrate at the synaptic membrane impairing the local tethering of lysosomes required to facilitate antigen extraction.

Given that Exo70 can also form complexes with BLOC-1, which through dysbindin and pallidin interaction regulates nuclear positioning (Monis et al, 2017; Ciciotte et al, 2003), we explored whether Exo70 was involved in nuclear positioning in B cells. For this purpose, we silenced Exo70 using a specific shRNA, and evaluated the localization of the nucleus with respect to the center of the immune synapse. As previously described (Sáez et al, 2019), our results show that Exo70-silenced B cells significantly decreased spreading areas compared to control cells (Fig. 6E,F). In addition, the nucleus of Exo70-silenced cells was displaced from the central region of the immune synapse (Fig. 6E,G). Additionally, we observed that the central pool of actin was more dispersed in the synaptic membrane (Fig. 6H,I) located in an excluded part of the poorly positioned nucleus. These results suggest that Exo70 has a role in nuclear positioning at the immune synapse, having displaced nuclei can result in a failure to position the lysosomes and therefore loss of lysosome secretion within precise domains of the immune synapse, required for an efficient antigen extraction.

Collectively, our results provide evidence that Nesprin-1 and Sun-1 regulate nuclear morphology and orientation towards the immune synapse of activated B cells. The connection between these nuclear envelope proteins, mainly though actin cytoskeleton, is required to orchestrate BCR internalization and promote precise lysosome recruitment for efficient antigen extraction.

## Discussion

We here show that B cells adjust their nuclear morphology upon interaction with immobilized antigens. Such changes involve the reorientation of the nuclear groove towards the antigen contact site (immune synapse) and changes in nuclear groove size, which rely mainly on interactions between nuclear envelope proteins, Nesprin-1 and Sun-1 (part of the LINC complex), and the actin cytoskeleton. Indeed, actin remodeling around the MTOC, at the perinuclear region, has been previously described to occur in B cells, where BCR engagement triggers actin depletion from MTOC in a proteasome-dependent manner, thereby enabling MTOC repositioning from the perinuclear region to the synaptic membrane (Ibañez-Vega et al, 2019). In this report, we reveal another level of regulation showing that the LINC complex connects the nucleus to actin in order to orientate and regulate the size of the nuclear groove facing the antigen at the immune synapse. In migrating fibroblasts, movement of the nucleus away from the leading edge, enables MTOC reorientation to the migrating front. Nuclear movement is associated to actin retrograde flow, a process regulated by Cdc42 (Gomes & Gundersen, 2006). Of note, Cdc42 is activated upon BCR antigen engagement, and regulates actomyosin contractions leading to MTOC, and lysosome polarization to the immune synapse (Yuseff et al, 2011a). Thus, Cdc42 emerges as a key candidate that could regulate nuclear morphology in B cells and shall now be evaluated.

The main role of MTOC polarization in B cells is to drive lysosome recruitment and exocytosis at the immune synapse to facilitate efficient antigen extraction, a crucial step for their activation (Yuseff et al, 2011a; Reversat et al, 2015). In this study, we propose that this process is also regulated by nuclear orientation and morphology where the nuclear groove becomes strategically positioned at the center of the synapse to enable the accumulation of lysosomes at the synaptic membrane.

What are the molecular links between nuclear positioning and lysosome trafficking? We recently showed that Exo70, a component of the exocyst complex, associates with the MTOC and is recruited to the synaptic membrane of B cells to promote lysosome tethering (Sáez et al, 2019). Interestingly, Exo70 has been described to interact with the lysosome-related organelle complex-1 (BLOC-1), which promotes Exo70 trafficking from the perinuclear region to the periphery of fibroblast. BLOC-1 also interacts with dysbindin and pallidin, which are involved in nuclear positioning (Monis et al, 2017). Knockout mice of dysbindin and pallidin, resulted in aberrant nuclei positioning in kidney tubule cells losing cell polarization (Ciciotte et al, 2003). In B cells, silencing Nesprin-1 and Sun-1 decreased the association of Exo70 with the MTOC, which displayed a more dispersed distribution throughout the synaptic membrane. Thus, proteins belonging to the LINC complex impact the distribution of proteins involved in lysosome positioning and tethering at the plasma membrane. Surprisingly, we observed that Exo70 is also required for correct nuclear positioning of the groove at the central region of the immune synapse. Whether Exo70 interacts with nuclear membrane proteins, such as the LINC complex, to regulate its positioning, remains to be addressed.

Interestingly, the internalization of the BCR was impaired in Nesprin-1 and Sun-1 silenced B cells, where the receptor remained arrested at the center of the immunological synapse. How do perturbations in nuclear envelope proteins, which result in nuclear mispositioning, affect the endocytosis of a cell surface receptor? One possibility is that Nesprin-1 and Sun-1 affect cortical actin organization, as shown here, which could compromise the recruitment of clathrins (Stoddart et al, 2005), actin binding proteins, such as Abp1, known to regulate BCR endocytosis (Onabajo et al, 2008) or by modulating the clustering of proteins coupled to this internalization (Hoogeboom & Tolar, 2015). How the nuclear envelope regulates cortical actin cytoskeleton organization and associated proteins involved in BCR endocytosis, remains to be addressed.

What other cues, in addition to BCR ligands, can modify nuclear morphology in B cells? Mechanical or physical forces that regulate intrinsic nuclear plasticity and morphology, have been described in other cells types. We noticed that the morphology of the plasma membrane frequently followed the form of the nucleus, which was more evident in primary B cells. Thus, mechanosensing at the level of the synaptic membrane could be directly transmitted to the nucleus. Studies in fibroblasts suggest that lamins A and C control the local mechanical stiffness within certain regions of the nuclear membrane, while lamin B, in turn mainly contributes to nuclear integrity (Lammerding et al, 2006). Most cells express both lamin A and B, and silencing of lamin A leads to irregular or deformed nuclei (Buchkovich et al, 2010; Adam & Goldman, 2012). Interestingly, B cells only express lamin B (Jansen et al, 1997), suggesting that their nuclei could be less rigid. In fact, we observed that, depending on the area of the antigen presenting surface, B cells expand their nuclear area, reflecting their high plasticity. This property would allow B cells to efficiently accommodate their cytoplasm to coordinate lysosome trafficking for antigen extraction and processing.

Additionally, nuclear plasticity could also regulate B cell motility inside the lymph node during the search for antigens. Indeed, a functional connection and coordination between the nucleus and the MTOC has been revealed in dendritic cells during antigen exploration. Migrating dendritic cells position their nucleus at the leading edge, which acts as a sensor to find the path of less resistance, whereas the MTOC defines directionality (Renkawitz et al, 2019). Whether B cells employ a similar strategy when searching for antigens, remains unknown. We did not observe any significant impairment in MTOC polarization to the immune synapse of B cells silenced for Nesprin-1 or Sun-1. However, given that in physiological environments B cells are frequently activated by antigens associated to softer substrates, it would be interesting to evaluate MTOC polarization in the absence of LINC complex components under such conditions.

Overall, this work highlights how the nucleus modifies its orientation and morphology in response to interaction with antigen-coated surfaces, thereby controlling intracellular reorganization and enabling B cells to efficiently organize immune synapses.

## Materials and Methods

### Antibodies and reagents described in supplementary materials and methods

#### Cell lines and culture

The mouse IgG+ B-lymphoma cell line IIA1.6 (Lankar et al, 2002) was used. Cells were cultured in CLICK medium (RPMI 1640 with 10% fetal bovine serum, Glutamax supplemented and 1 mM sodium pyruvate, 100 µg/ml streptomycin, 100 U/ml penicillin and 0.1% β-mercaptoethanol (Vascotto et al, 2007). HEK 293T cells were cultured for lentiviral production in DMEM supplemented with 10% FBS and penicillin/streptomycin. Cell culture products were purchased from Life Technologies. Spleen derived primary B cells were isolated from C57BL/6 mice using a magnetic cell sorting B cell isolation kit (Miltenyi) according to the manufacturer’s instructions. Mice protocols were approved by the Institutional Scientific Ethics Committees for Animal and Environmental Care and Research Biosafety, Pontificia Universidad Católica de Chile.

#### Cell transfection

The Amaxa Cell Line Nucleo-factor Kit R (L-013 program; Lonza) was used to electroporate 5 × 10^6^ IIA1.6 B Lymphoma cells for different transfections. For Centrin-GFP plasmid transfection we used 2 µg of DNA, and cells were cultured for 16 h before functional analysis. In the case of transfection with siRNA we used Silencer® Select (Ambion/Thermofisher) against Nesprin-1(Syne1): s234287 (#1), s234288 (#2); siRNA Sun1: s94911 (#1), s94912 cells (#2), these were incubated for 48 h before analysis. As a control, we used a scrambled siRNA (Qiagen) at 10nM.

#### Antigen-coated beads and Antigen-coated dishes

We used two assays to activate B cells; Antigen-coated beads and Antigen-coated dishes prepared as previously described (Yuseff et al, 2011b). To evaluate nuclear orientation, nuclear groove area and polarization events, ∼2 × 10^7^ 3-μm latex NH2-beads (Polyscience) were activated with 8% glutaraldehyde for 4 h at room temperature. Beads were washed with cold PBS and incubated overnight at 4°C with 100 μg/mL of F(ab’)2 goat anti-mouse immunoglobulin G (IgG) (MP Biomedical), referred to as Antigen or activating beads. For antigen extraction assays, beads were coated with IgG plus OVA 100 μg/mL. To evaluate nuclear rotation and immune synapse organization we performed antigen-coated dishes assays, where 10 micron glass dishes were coated with IgG in PBS overnight at 4°C.

#### Activation of B cells and immunofluorescence

Cells were plated on poly-L-Lysine– coated glass coverslips and activated with Ag-coated beads (1:1 ratio) for different time points at 37°C, 5% CO2, and then fixed in 4% paraformaldehyde (PFA) for 10 min at room temperature. Fixed cells were incubated with antibodies according to their manufacturer’s instructions. Time-Lapse of nucleus and MTOC rotations protocol is indicated in Supplementary Materials and Methods.

#### Ag extraction assay

For antigen extraction assays were performed as previously described (Yuseff et al, 2011b). B cells incubated in a 1:1 ratio with IgG-OVA-coated beads were plated on poly-Lys cover-slides at 37°C 5% CO2, fixed and stained for OVA. The amount of OVA remaining on the beads was calculated by establishing a fixed area around beads in contact with cells and measuring fluorescence on three-dimensional (3D) projections obtained from the sum of each plane.

#### Cytoplasm, nuclear and perinuclear fractioning

Subcellular fractioning was performed as previously described for (Shaiken & Opekun, 2014), that we adapted to B cells. For protocol descriptions, see supplementary Methods.

#### Immunoprecipitation assay

Immunoprecipitation was performed as previously described by (Zhang et al, 2005), adapted to B cells. For protocol descriptions, see supplementary Methods.

#### Cell imaging

For all confocal microscopy experiments images were obtained with a Zeiss LSM880 Airyscan Confocal microscope with a 63X/1.4NA oil immersion lens, in 16-bit Z-stack image sets with 0.2 µm optical section and 0.07 µm x 0.07 µm pixel size. For OVA extraction assays, epifluorescence imaging was used. Z-stack images were obtained with 0.5 microns between slices. Images were acquired in an epifluorescence microscope (Nikon Ti2Eclipse) with a 60 X/1.25NA oil immersion objective.

### Image analysis

Image processing and analysis were performed with FIJI/ImageJ software (Schindelin et al, 2012). 3D analysis was performed using the SCIAN-Soft software tools (https://github.com/scianlab), programmed on the IDL programming language (ITT/Harris Geospatial; Boulder, CO). Image brightness and contrast were manually adjusted for visualization purposes but not for analysis (e.g. in the case of segmentations).

– **Nuclear groove area and reorientation**: Nuclear groove area was calculated by manually delineating the principal groove at its maximum size (from staining with lamin B). Reorientation was calculated by measuring the angle between the centers of mass of the bead (a), cell (b), and nuclear groove (c) (Fig. 1B). We then calculated the percentage of cells with the nuclear groove: oriented toward the bead (Polarized) with the angle between 0° and 45°; centered nuclear groove, with the angle was between 45° and 90° (Centr P) or 90-135° (Centr AP), and antipolarized with the angle was between 135 and 180°.
– **MTOC and LAMP polarity indexes** were determined as previously described (Ibañez-Vega et al, 2019). Briefly, we manually selected the location of the MTOC and delimited the cell and we also obtained the center of mass of the cell, (Cellmc) and the bead (Beadmc). The position of the MTOC was projected on the vector defined by Cellmc and Beadmc axis (Pj-mtoc). The MTOC polarity index was calculated by dividing the distance between Cellmc and Pmtoc and the distance between Cellmc and Beadmc. The index ranged from -1 (anti-polarized) to +1 (fully polarized). LAMP polarity index was calculated using the center of mass of the LAMP-1 staining and we continue the analysis as the MTOC polarity index). Lysosomes at the IS in the beads model (lysosome rings phenotype), was quantified by measuring the LAMP1 fluorescence intensity in the circular area closely surrounding the bead (3.5 μm) and normalized by total LAMP1 fluorescence intensity.
– **Time-Lapse of nucleus and MTOC rotations**: MTOC was labeled with Centrin-GFP and nucleus with Hoechst. Single-cell images shown in the figures were cropped from a larger field.
– **Nuclear or actin volume**: Confocal image stacks were first filtered in FIJI using the Trainable Weka Segmentation plugin (Arganda-Carreras et al, 2017), to enhance membrane signals. Next, 2D threshold filters were applied within SCIAN-Soft in order to produce binary images of the nuclear/actin fluorescent signal as regions of interest (ROIs). **Actin surrounding the nucleus**: Nuclear actin ROIs were defined by a logical filter to obtain the intersection between the actin ROIs and the nucleus ROI. In each case (nucleus, actin, and nuclear-actin), adjacent ROIs along the z axis were connected to define 3D ROIs, and their volume was quantified by voxel counting.
– **Nuclear rotation in IgG coated dishes**: The rotation of the nucleus in covers was measured by taking the angle left by the baseline between the two main lobes, which surround the nuclear groove, at its maximum area (Fig. 3E). The angle is an analysis taken from the FIJI program.
– **The z radial distribution of lysosomes or BCR** was calculated by measuring their mean fluorescence intensity (MFI) in the cell for each z-slice (0.2 µm) from the IS until 2 µm inside the cell (10 z-slices). Next, the MFI values were normalized by dividing each one for the total fluorescence of these slices, and multiplied them by 100.
– **Segmentation and position of lysosome clusters relative to the IS**. Accumulation of lysosomes at the IS was quantified by measuring the LAMP1 fluorescence intensity. Using a custom made FIJI macro, lysosome clusters formation was quantified by calculating their area inside and outside the immune synapse and nucleus region. The procedure was defined as follows. Cell (the limits of actin) and nucleus areas were manually outlined within a reference z-slice (immune synapse). Lysosome clusters positioning with respect to the IS was determined by measuring the LAMP1 MFI in 1 μm decreasing steps, from the nucleus perimeter inwards; and in 1 μm increasing steps from the nucleus perimeter to the cell boundary. LAMP1 MFI within the nucleus area was considered as lysosomes clusters being at the immune synapse, whereas the fluorescence signal outside the nuclear area was considered as lysosomes clusters displaced from the immune synapse.
– **Distribution of Exo70**: The xy radial distribution of exo70 in resting and activated B cells was calculated with the “Radial Profile” FIJI plugin available at https://imagej.nih.gov/ij/plugins/radial-profile.html. By setting a reference point for the MTOC, MFI was calculated and plotted for circular regions for a growing radius, from 0 to 2 µm.

### Statistical analysis

Statistical analysis was performed with GraphPad Prism (GraphPad Software; San Diego, CA). Shapiro-Wilk test was previously used to contrast the normality of the data set and thus decide which analytical test will be used (parametric or nonparametric). The p values were calculated using different tests, as indicated in figure legends.

## Acknowledgements

We thanks M. Bornens for the eGFP-centrin1 plasmid. This project was supported by a research grant from FONDECYT, JD (#1171024) and MI-Y (#1180900). We are grateful to Unidad de Microscopía Avanzada (UMA), Pontificia Universidad Católica de Chile, specialy to Nicole Salgado and Fernanda Garate for their support in images acquisition. Felipe Del Valle and Martina Alamo for critical feedback and ideas.

## Author contributions

RU performed most of the experiments. OC, FC performed experiments and helped draft the manuscript, JJ provided tools and feedback for 2D and 3D analysis. JL helped with image analysis, JJ-S performed co-immunoprecipitation and perinuclear fractionation assays, CR performed immunofluorescence and 3D analysis. ERG, CC, SH provided tools for analysis and development of the project. MI-Y and JD provided tools and critical feedback on experiments and write the manuscript. JD also performed experiments, analysis and conceived and supervised the project.

---

**Figure EV1.**
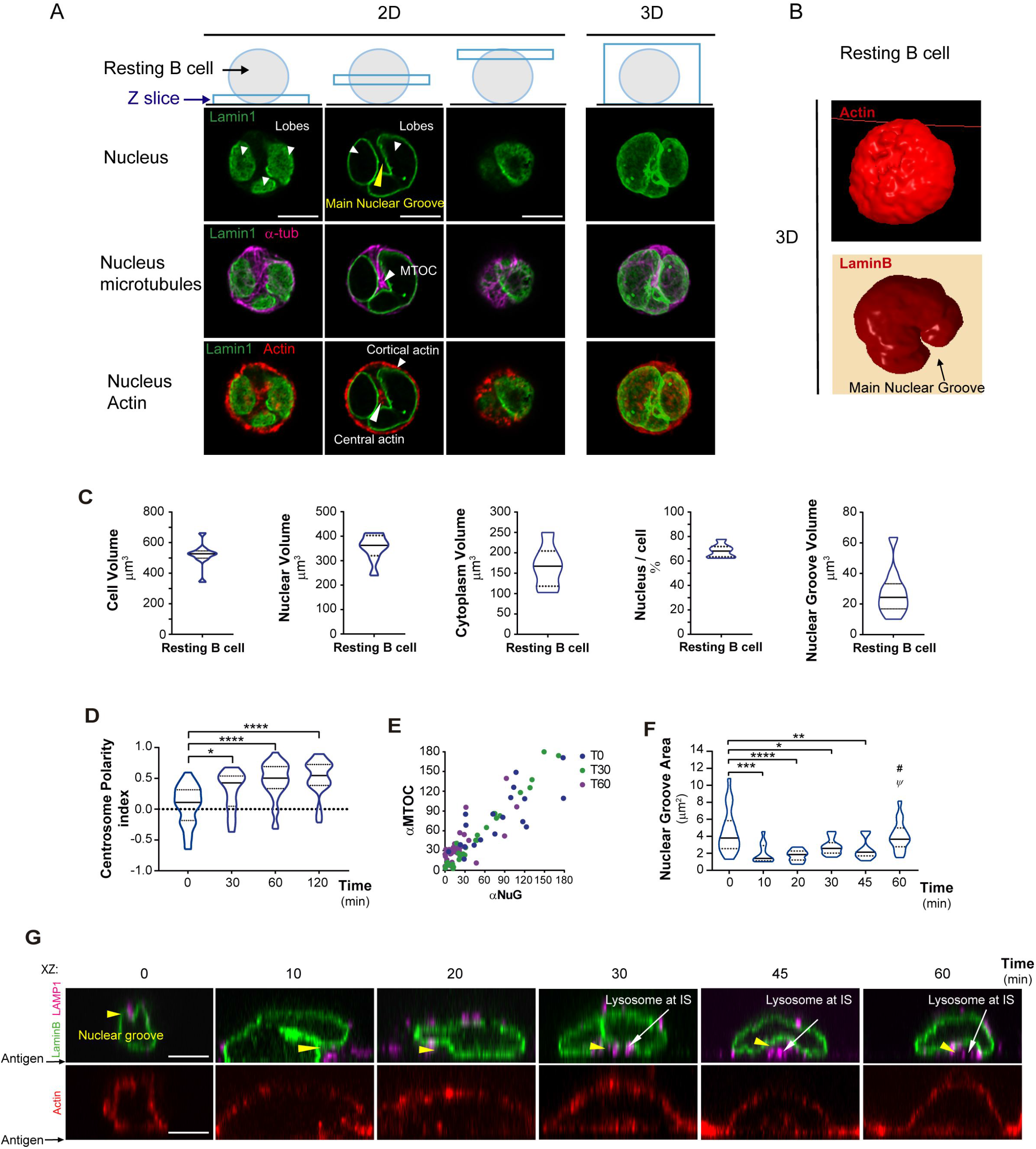

**Figure EV2.**
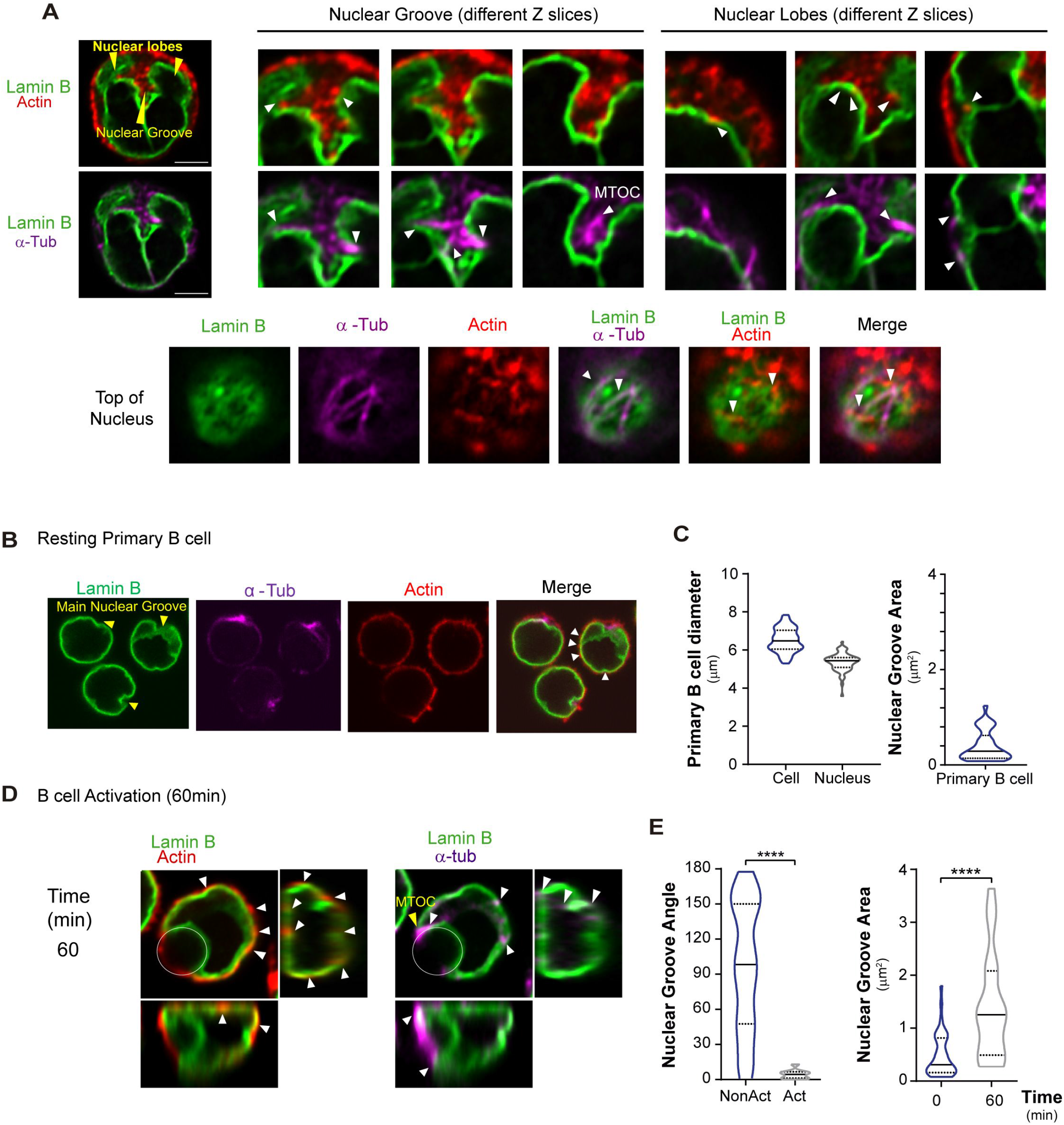

**Figure EV3.**
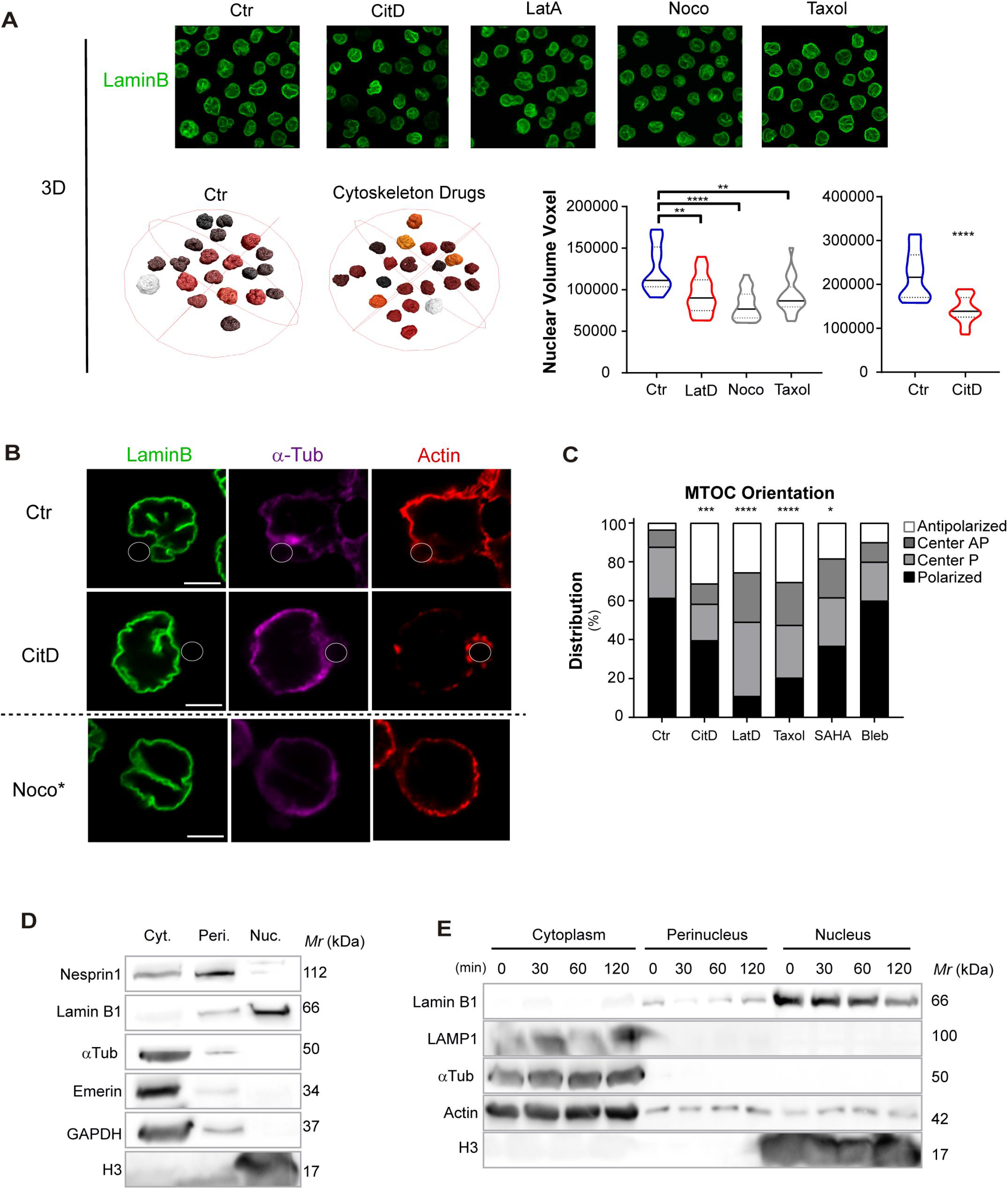

**Figure EV4.**
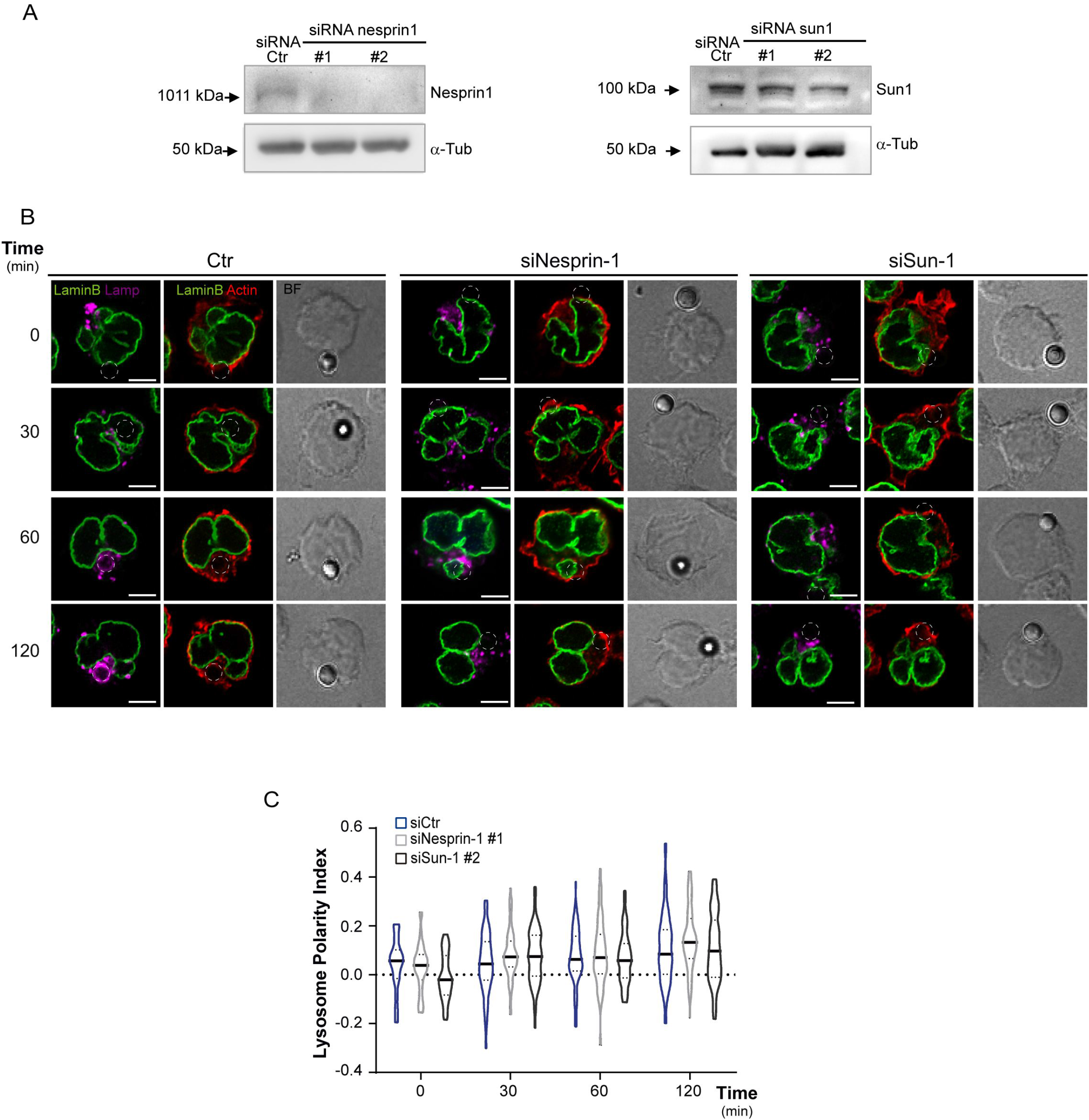

**Figure EV5.**
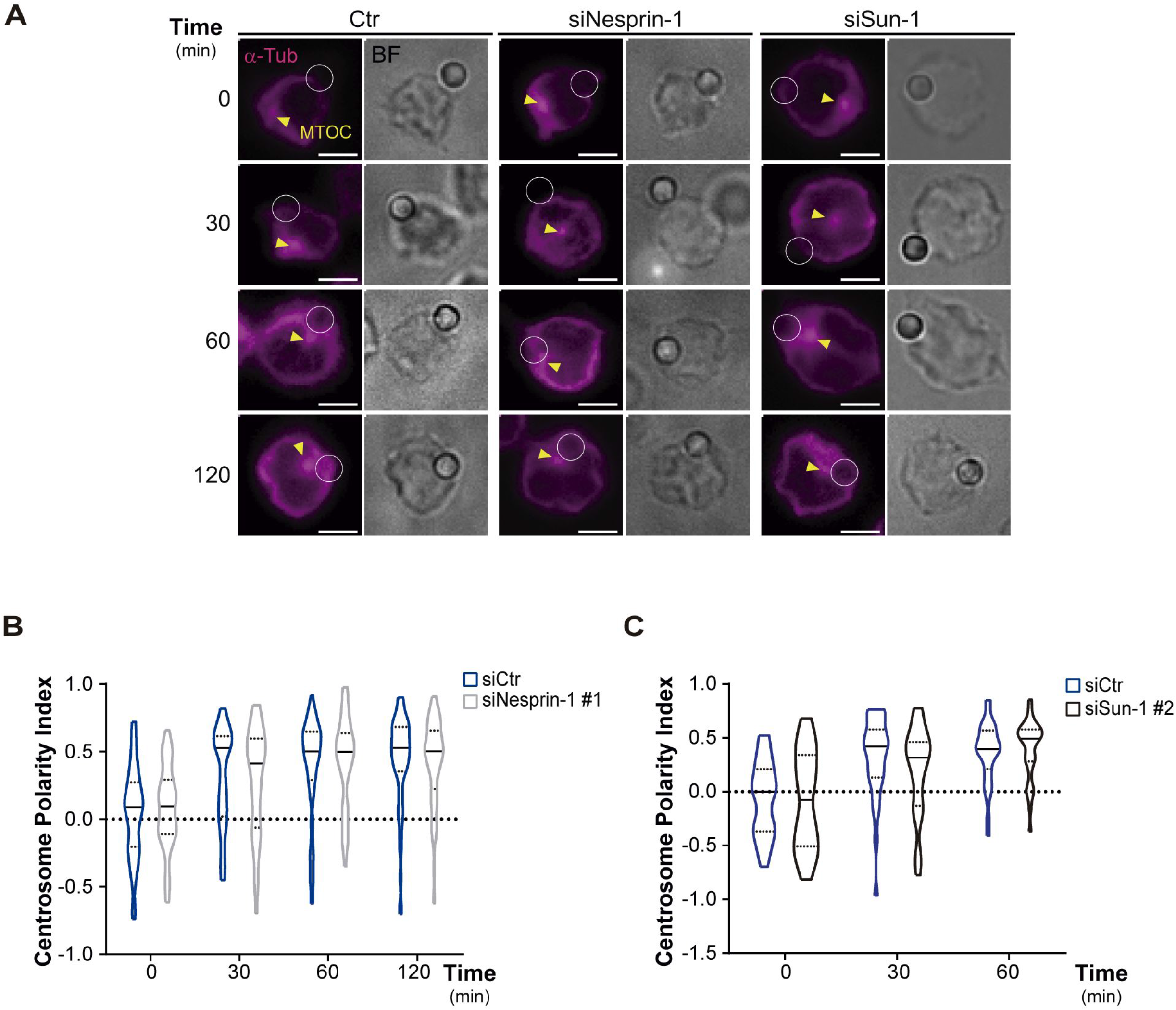

**Figure EV6.**
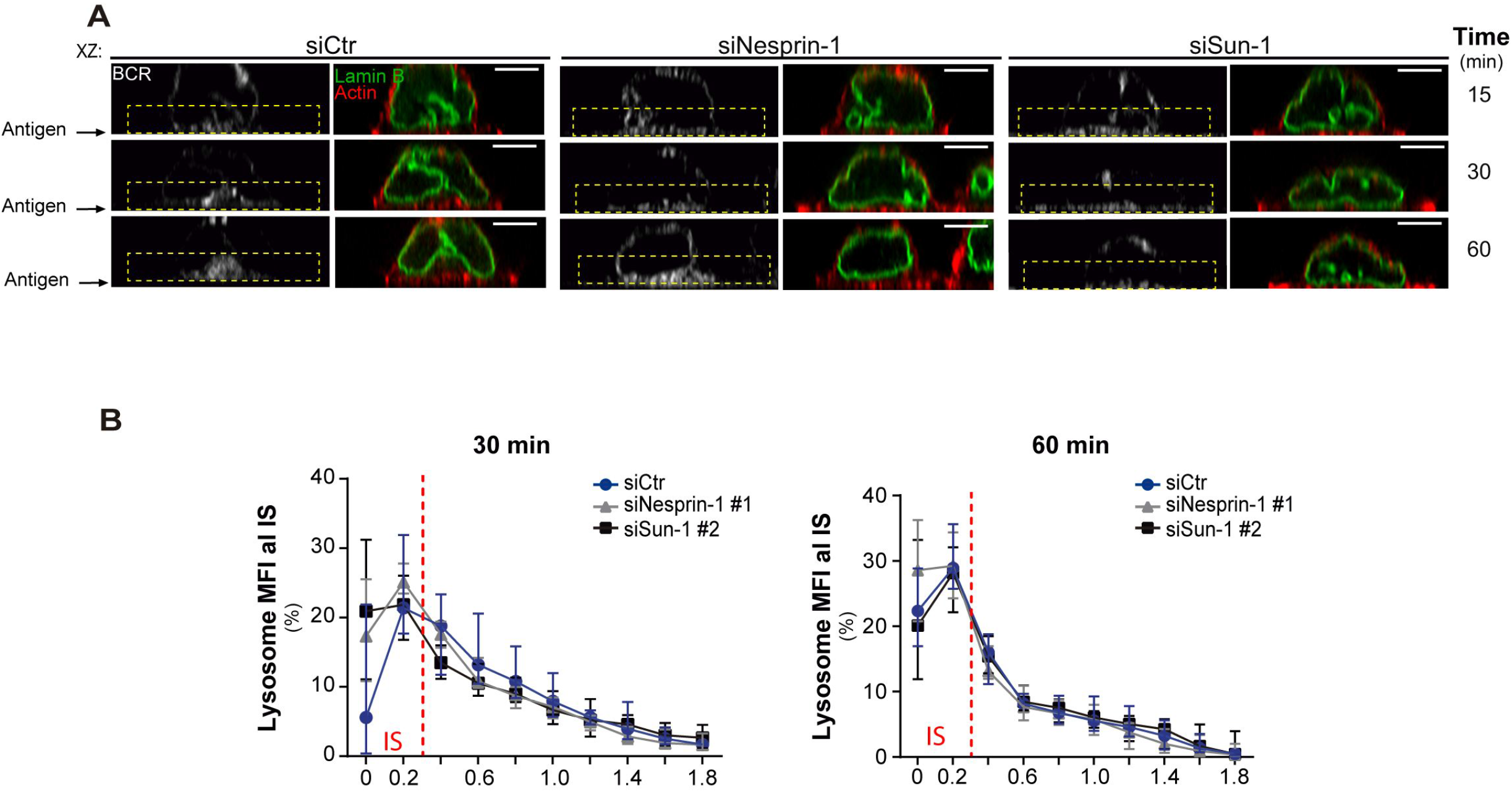

